# Deletion of the transcriptional regulator TFAP4 accelerates c-MYC-driven lymphomagenesis

**DOI:** 10.1101/2022.12.19.520971

**Authors:** Margaret A Potts, Shinsuke Mizutani, Alexandra L Garnham, Connie S N Li Wai Suen, Andrew J Kueh, Lin Tai, Martin Pal, Andreas Strasser, Marco J Herold

## Abstract

Many lymphoid malignancies arise from deregulated c-MYC expression in cooperation with additional genetic lesions. While many of these cooperative genetic lesions have been discovered, DNA sequence data suggest that many more do exist. However, their contributions to c-MYC driven lymphomagenesis have not yet been investigated. We identified TFAP4 as a potent suppressor of c-MYC driven lymphoma development in a previous genome-wide CRISPR knockout screen in primary cells *in vivo* (*Mizutani et al, 2022*). CRISPR deletion of TFAP4 in *Eμ-MYC* transgenic hematopoietic stem and progenitor cells (HSPCs) significantly accelerated c-MYC-driven lymphoma development in mice. TFAP4 deficient *Eμ-MYC* lymphomas all arose at the pre-B cell stage. Characterization of the transcriptional profile of pre-leukemic pre-B cells in *Eμ-MYC/Cas9/sgTFAP4* transplanted mice, revealed that TFAP4 deletion reduced expression of several master regulators of B cell differentiation, such as *Spi1, SpiB* and *Pax5*, which all have been shown to be bound by TFAP4. We therefore conclude that loss of TFAP4 leads to a block in differentiation during early B cell development, causing accelerated c-MYC-driven lymphoma development.

## Introduction

The oncogene *c-MYC* is abnormally highly expressed *in* ~70% of human cancers (Dang, 2012). Notably, c-MYC is commonly overexpressed in hematological malignancies, such as Burkitt lymphoma (BL), a B cell malignancy driven by the t(8;14) chromosomal translocation that subjugates the *c-MYC* gene under the control of the immunoglobulin heavy (*IgH*) chain gene enhancer *Eμ* (Haluska *et al*, 1986). The *Eμ-MYC* transgenic mouse serves as a pre-clinical model of BL and other malignancies driven by deregulated expression of c-MYC. These mice constitutively express human c-MYC under the control of the murine *Eμ* enhancer throughout B cell development leading to increased cycling and expansion of early B cell progenitors that subsequently transform into disseminated lymphoma following the acquisition of cooperating oncogenic mutations (Adams *et al*, 1985; Harris *et al*, 1988; Langdon *et al*, 1986).

Since the development of BL and other lymphoid malignancies requires additional genetic and epigenetic aberrations that cooperate with deregulated c-MYC expression, we previously set out to identify such cooperating mutations in an unbiased genome-wide functional genetic screen *in vivo* (Mizutani *et al*., 2022). From this screen, we identified the transcription factor TFAP4 (also known as AP-4) as one of the top hits (Mizutani *et al*., 2022). TFAP4 is a basic helix-loop-helix leucine zipper transcription factor reported to both activate or repress target gene expression upon homo-dimerization (Mermod *et al*, 1988). *Tfap4* expression itself is directly regulated by c-MYC. TFAP4 contributes to several c-MYC regulated cellular processes, including cell proliferation, cell cycle control and cellular senescence, by functional interaction with c-MYC including binding to the same gene promoter elements (reviewed by Wong *et al*., (Wong *et al*, 2021)). Of note, TFAP4 is critical for c-MYC-driven proliferation and maturation of B cells in germinal centers following an infectious challenge (Chou *et al*, 2016). This reveals that TFAP4 is an important transcription factor acting downstream of and/or in parallel with c-MYC to regulate the proliferation and Ig class switching of mature B lymphoid cells.

In cancer, TFAP4 has predominantly been identified as an oncogene that has been implicated in driving tumorigenesis by impacting the epithelial to mesenchymal transition, “stemness” and several other cellular processes (Wong *et al*., 2021). In some human cancers high levels of TFAP4 expression correlates with poor prognostic outcomes (Hu *et al*, 2013; Wei *et al*, 2017). However, these oncogenic properties of TFAP4 were found in solid cancers, such as breast cancer (Chen & Chiu, 2015)’, colorectal carcinoma (Jaeckel *et al*, 2018; Shi *et al*, 2014), gastric cancer (Liu *et al*, 2012), neuroblastoma (Boboila *et al*, 2018; Xue *et al*, 2016) and others (Chen *et al*, 2017; Huang *et al*, 2019), and the role of TFAP4 in hematological malignancies has only been investigated in one study to date (Tonc *et al*, 2021).

Several bioinformatic studies have analyzed gene expression and mutation status in BL human patient samples with the aim of further classifying somatic mutations as driver genes that cooperate with c-MYC in malignant transformation (Grande *et al*, 2019; Kaymaz *et al*, 2017; López *et al*, 2019). While frequently mutated genes and altered pathways in BL, such as the TP53 and PI3K regulated pathways, have been well characterized, there are less commonly, but still significantly, mutated genes, whose contributions to c-MYC-driven lymphomagenesis are not well understood. Interestingly, mutations in *TFAP4* were identified in 9.7% of BL patient samples (Grande *et al*., 2019). Of note, and conversely to solid cancers (see above), these were predominantly inactivating mutations, suggesting TFAP4 has tumour suppressive functions in this context. However, this type of analysis does not provide mechanistic insights into how defects in TFAP4 contribute to tumorigenesis.

We employed *Eμ-MYC/Cas9* doubly transgenic mice and a fetal liver HSPC transplantation approach to investigate the tumour suppressive role of TFAP4 during c-MYC-driven lymphoma development. We found that loss of TFAP4 led to malignant transformation at the pre-B cell developmental stage, prior to surface Ig (sIg) expression. Gene expression analysis of pre-leukemic pre-B cells revealed that several transcription factors that are critical for B cell differentiation, and which are also direct targets of TFAP4, were abnormally reduced when TFAP4 was absent. This demonstrates that TFAP4 binds to and regulates key genes responsible for B cell differentiation and maturation and in *Eμ-MYC* transgenic B lymphoid cells its loss dysregulates differentiation resulting in expansion of the rapidly proliferating pre-B cell pool that transforms into fully malignant c-MYC driven lymphoma.

## Results

### Deletion of TFAP4 in HSPCs accelerates c-MYC-driven lymphomagenesis

We set out to validate *Tfap4* as a top hit accelerating *Eμ-MYC* induced lymphomagenesis identified from a previous genome-wide CRISPR knockout screen (Mizutani *et al*., 2022) using independent sgRNAs targeting *Tfap4*. To this end, we isolated HSPCs from fetal livers of *Eμ-MYC/Cas9* doubly transgenic day 13.5 embryos (E13.5) on a C57BL/6-Ly5.2 background and independently transduced them with two sgRNAs targeting *Tfap4* (*sgTfap4*), a positive control sgRNA targeting *Trp53 (sgTrp53*) (Aubrey *et al*, 2015) or a non-targeting negative control sgRNA targeting human *BIM* or human *NLRC5 (sgControl*). Subsequently, the sgRNA transduced *Eμ-MYC /Cas9* HSPCs were transplanted into lethally irradiated C57BL/6-Ly5.1 recipient mice that were then monitored for lymphoma development (Figure 1a).

**Figure 1.**
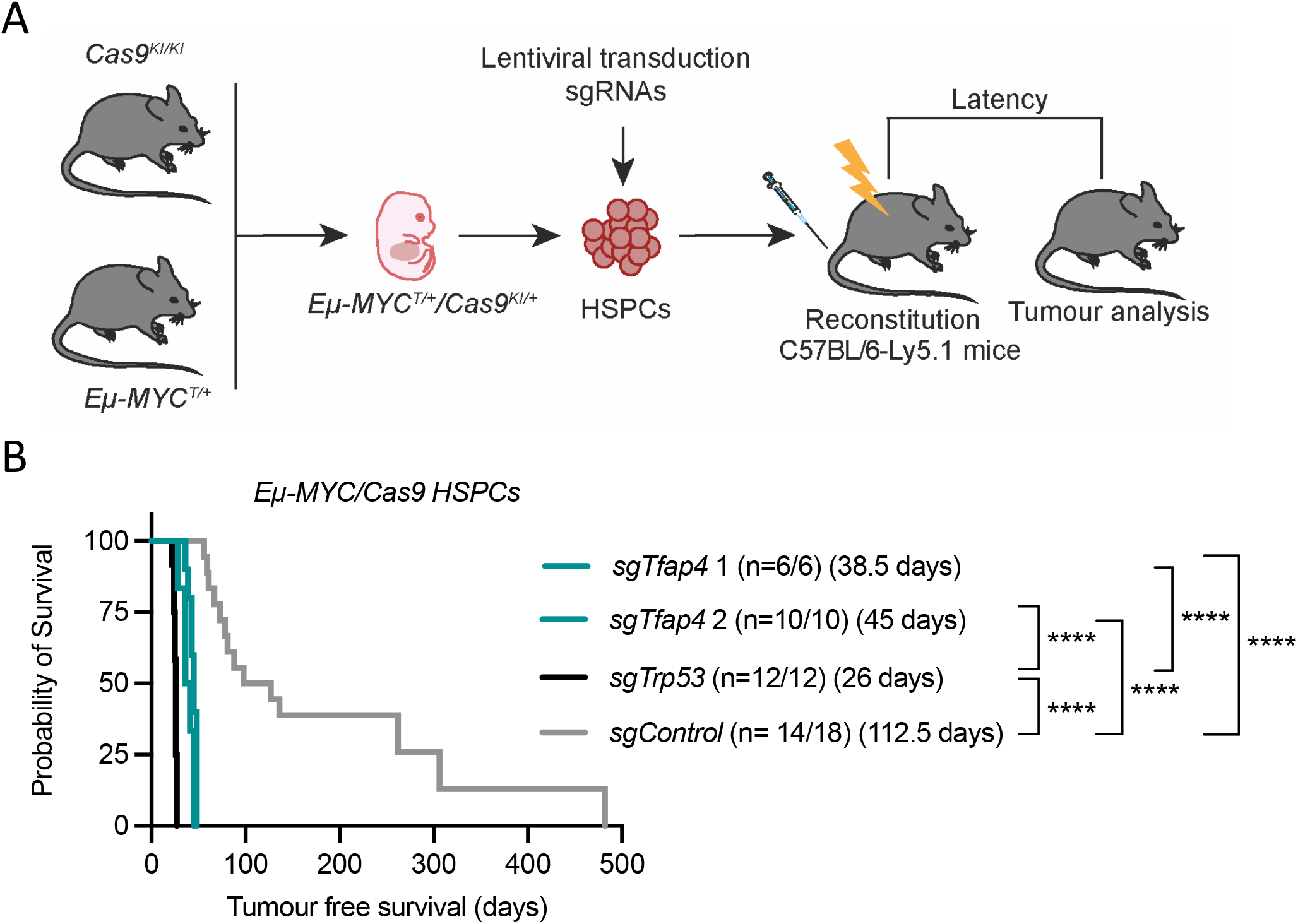
Loss of *Tfap4* accelerates c-MYC-driven lymphomagenesis. **A,** Schematic representation of hematopoietic reconstitution of lethally irradiated recipient mice with fetal liver derived donor HSPCs. Fetal liver cells from double transgenic *Eμ-MYC/Cas9* E13.5 donor embryos (an abundant source of HSPCs) were lentivirally transduced with vectors encoding BFP or CFP as a marker and sgRNAs targeting *Tfap4 (sgTfap4*), a positive sgRNA targeting *Trp53 (sgTrp53*) or a negative non-targeting control sgRNA (*sgControl*). These transduced donor fetal liver cells were then transplanted into lethally irradiated C57BL/6-Ly5.1 recipient mice. Tumour-free survival of recipient mice was measured as days post-transplantation. Hematopoietic tissues and peripheral blood were harvested from tumour burdened recipient mice for further analysis. **B,** Kaplan-Meier survival curve showing tumour-free survival of mice transplanted with either of two vectors containing a *sgTfap4*, a positive control *sgTrp53* or a negative *sgControl*. N indicates number of sick mice/number of mice transplanted with sgRNA transduced HSPCs for each sgRNA. Median survival post transplantation in days is indicated in brackets. \*\*\*\**P* < 0.0001.

All recipient mice progressed to display characteristic *Eμ-MYC* lymphoma pathology, including enlarged lymph nodes, spleen and thymus, and elevated white blood cell count (WBC) as well as thrombocytopenia (Supplementary Figure 1a, b). Mice transplanted with the *sgTfap4/Eμ-MYC/Cas9* HSPCs developed lymphoma at a slightly slower pace than mice that had been transplanted with *sgTrp53* transduced *Eμ-MYC/Cas9* HSPCs (median survival of *sgTfap4-1* = 38.8 days; *sgTfap4-2* = 45.5 days *vs sgTrp53* = 26 days), but significantly faster than mice transplanted with *sgControl* transduced *Eμ-MYC/Cas9* HSPCs (112.5 days) (Figure 1b). This finding validates that deletion of *Tfap4* markedly accelerates c-MYC-driven lymphomagenesis.

### The absence of TFAP4 did not change reliance on pro-survival BCL-2 family members to accelerate MYC-driven lymphomagenesis

Avoiding cell death is one of the hallmarks of cancer (Hanahan, 2022). This can be achieved by abnormally increased expression of pro-survival BCL-2 family members (Strasser *et al*, 1990). *Eμ-MYC* lymphoma cells are generally dependent on the pro-survival BCL-2 family member MCL-1 for their sustained survival (Campbell *et al*, 2010; Kelly *et al*, 2014). To assess whether loss of *Tfap4* changes the dependency of *Eμ-MYC* lymphoma cells on pro-survival BCL-2 family members, we generated cell lines from the primary *sgTfap4/Eμ-MYC/Cas9*, in which TFAP4 protein was absent (Figure 2a), and *sgControl*/*Eμ-MYC/Cas9* lymphomas, which express TFAP4 protein (Figure 2a), and examined their responses to the BH3 mimetic drugs ABT-199/Venetoclax (Roberts *et al*, 2016) or S63845 (Kotschy *et al*, 2016) that inhibit BCL-2 or MCL-1, respectively. No differences in cell viability were observed between the *sgTfap4 vs sgControl/Eμ-MYC/Cas9* treated cells as measured at 24 h of treatment with these agents by flow cytometric analysis (Supplementary Figures 2a, b, c, f, g). These findings demonstrate that the absence of TFAP4 accelerates c-MYC-driven lymphoma development through mechanisms other than altering the dependency of these malignant cells on select pro-survival BCL-2 proteins.

**Figure 2.**
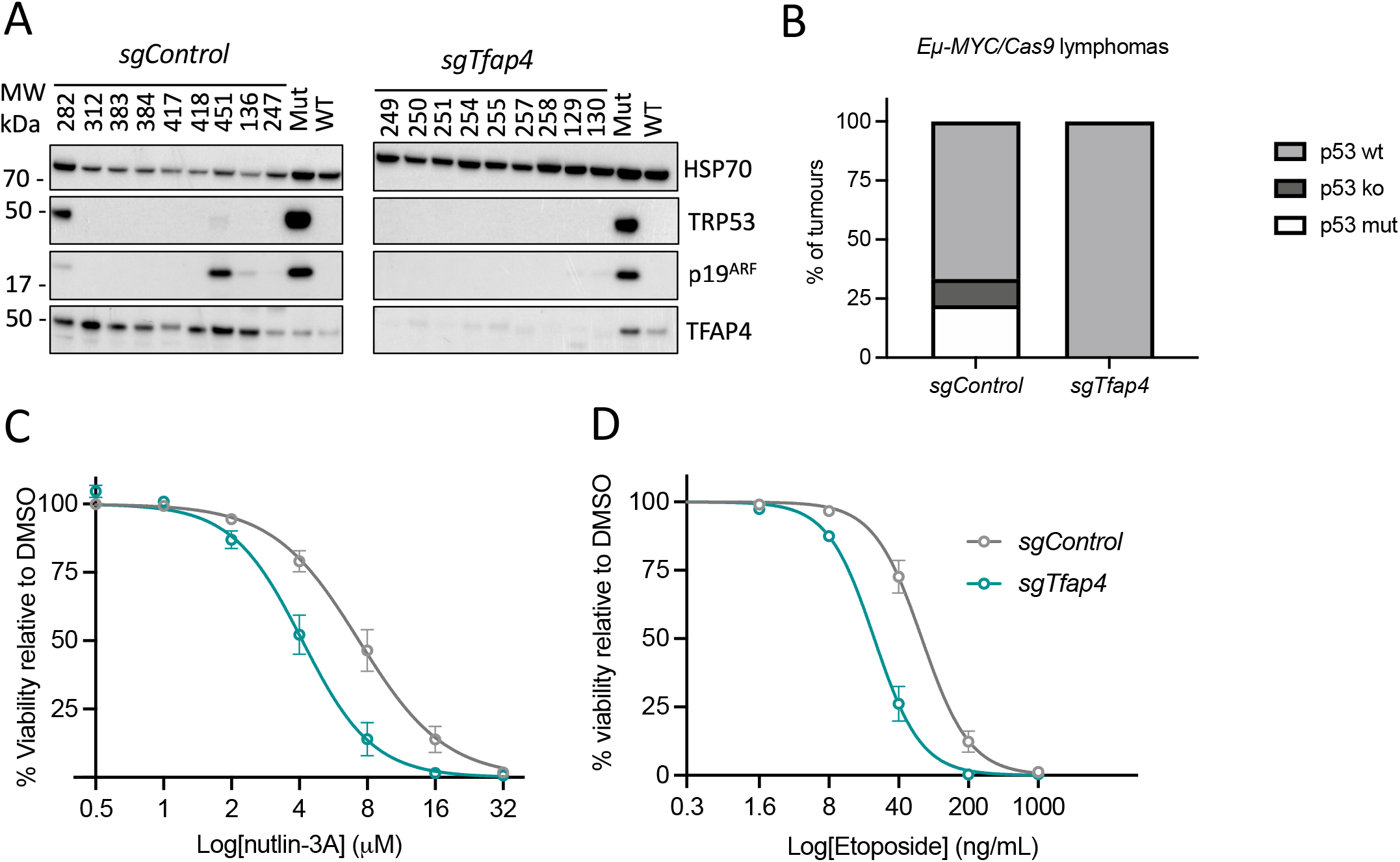
Deletion of *Tfap4* removes the selection pressure to acquire defects in the TRP53 pathway during c-MYC-driven lymphoma development. **A,** Western blot analysis of *sgTfap4/Eμ-MYC/Cas9* and *sgControl/Eμ-MYC/Cas9* primary lymphomas for TRP53, p19/ARF, TFAP4 and HSP70 (protein loading control). Includes positive and negative control cell lysates for mutant (Mut) and wild-type (WT) TRP53. Protein weights are indicated in kDa. **B,** Summary graph representing percentages of *sgTfap4/Eμ-MYC/Cas9* (n=9) and *sgControl/Eμ-MYC/Cas9* (n=9) lymphomas that have defects in the TRP53 pathway as assessed by Western blotting for TRP53 and p19/ARF. TRP53 wt determined as lack of detectable protein expression of both TRP53 and p19/ARF. TRP53 knockout (ko) is determined as no detectable TRP53 protein expression with high expression of p19/ARF. Mutant TRP53 is identified by high level expression of both TRP53 and p19/ARF proteins. **C-D,** Cell viability response curves in *sgTfap4/Eμ-MYC/Cas9* and *sgControl/Eμ-MYC/Cas9* lymphoma cell lines 24 h after treatment with the indicated doses of nutlin-3A **(C)** or etoposide **(D)**. Cell viability was determined by flow cytometry; live cells were identified as the AnnexinV/PI double negative population. Data represent percentage mean survival of lymphoma cell lines at each dose (*sgControl* n=3, *sgTfap4* n=4-9). Data were log transformed and fitted to non-linear regression mean ±SEM.

### TFAP4 deficient *Eμ-MYC* lymphomas are not selected for defects in the TRP53 pathway

Since approximately 25% of lymphomas that spontaneously develop in *Eμ-MYC* transgenic mice show defects in the TRP53 pathway, we examined the *sgTfap4/Eμ-MYC/Cas9* lymphomas for abnormalities in this pathway. Interestingly, no TRP53 pathway alterations were detected in any of the *sgTfap4/Eμ-MYC/Cas9* lymphoma cells tested by Western blotting for the levels of TRP53 and p19/ARF proteins (high levels of these proteins are an indicator for mutations in the *Trp53* gene and/or TRP53 pathway defects (Figure 2a, b) (Eischen *et al*, 1999)). This contrasts with the *sgControl/Eμ-MYC/Cas9* lymphoma cell lines, with ~30% of those exhibiting TRP53 protein/pathway aberrations. Functionality of the TRP53 pathway was confirmed in *sgTfap4*/*Eμ-MYC/Cas9* lymphoma cells lines by showing that they are readily killed by treatment with nutlin-3A, a drug that blocks MDM2, the E3 ubiquitin ligase that targets TRP53 for proteasomal degradation (Vassilev *et al*, 2004) (Figure 2c, Supplementary Figure 2d) and the DNA-damaging chemotherapeutic drug etoposide (Figure 2d, Supplementary Figure 2e) that can also kill lymphoma cells through activation of TRP53 (Strasser *et al*, 1994). Collectively, these data reveal that loss of TFAP4 reduces the selection pressure for *Eμ-MYC* lymphomas to acquire defects in the TRP53 tumour suppressor pathway.

### TFAP4 impairs differentiation of *Eμ-MYC* transgenic B lymphoid cells at the pre-B stage

Interestingly, we found that *sgTfap4/Eμ-MYC/Cas9* lymphoma cell lines all represented neoplastic counterparts of surface Ig negative pre-B cells. In contrast, the *sgControl* as well as *sgTrp53 Eμ-MYC/Cas9* lymphomas comprised a mixture of sIg negative pre-B lymphomas and sIg positive B cell lymphomas (Figure 3a, b). This suggests that the absence of TFAP4 drives transformation at an early stage of B cell development. To better understand how TFAP4 deletion contributes to c-MYC-driven lymphoma development, a pre-leukemic analysis was performed. Lethally irradiated wt (C57BL/6-Ly5.1) recipient mice were transplanted with either *sgTfap4* or *sgControl* transduced *Eμ-MYC/Cas9* HSPCs, but this time hematopoietic cells of these recipient mice were harvested three weeks post-transplantation, a time point early during lymphomagenesis when their B lymphoid cells are not yet fully transformed. Bone marrow, spleen, thymus, lymph nodes and peripheral blood were analyzed by flow cytometry to determine the proportions of the various B as well as T lymphocyte and myeloid cell subsets. No differences were observed in the total cellularity of the hematopoietic tissues between the *sgTfap4/Eμ-MYC/Cas9 vs* the *sgControl/Eμ-MYC/Cas9* groups (Supplementary Figure 3a). Furthermore, the contribution of donor cells carrying both *Cas9* (GFP positive) and the sgRNA (BFP positive) was comparable between the *sgTfap4/Eμ-MYC/Cas9 vs* the *sgControl/Eμ-MYC/Cas9* HSPCs transplanted cohorts across all hematopoietic tissues examined (Supplementary Figure 3b). However, these compartments were not completely composed of donor HSPC derived cells. This can be explained because: (1) not all *Cas9-GFP* donor cells carried *sgRNA-BFP* due to the infection efficiency of the donor HSPCs; (2) not all host cells have been cleared from the hematopoietic tissues of recipient mice examined three weeks post-transplantation with donor cells. The latter is especially apparent for T lymphocytes and granulocytes: only a small percentage of these cell types were donor derived (Supplementary Figure 3 c, d). Thus, to assess the impact of *Tfap4* deletion in different cell types, only sgRNA/Cas9 double expressing (BFP^+^ GFP^+^) *Eμ-MYC* cells were examined.

**Figure 3.**
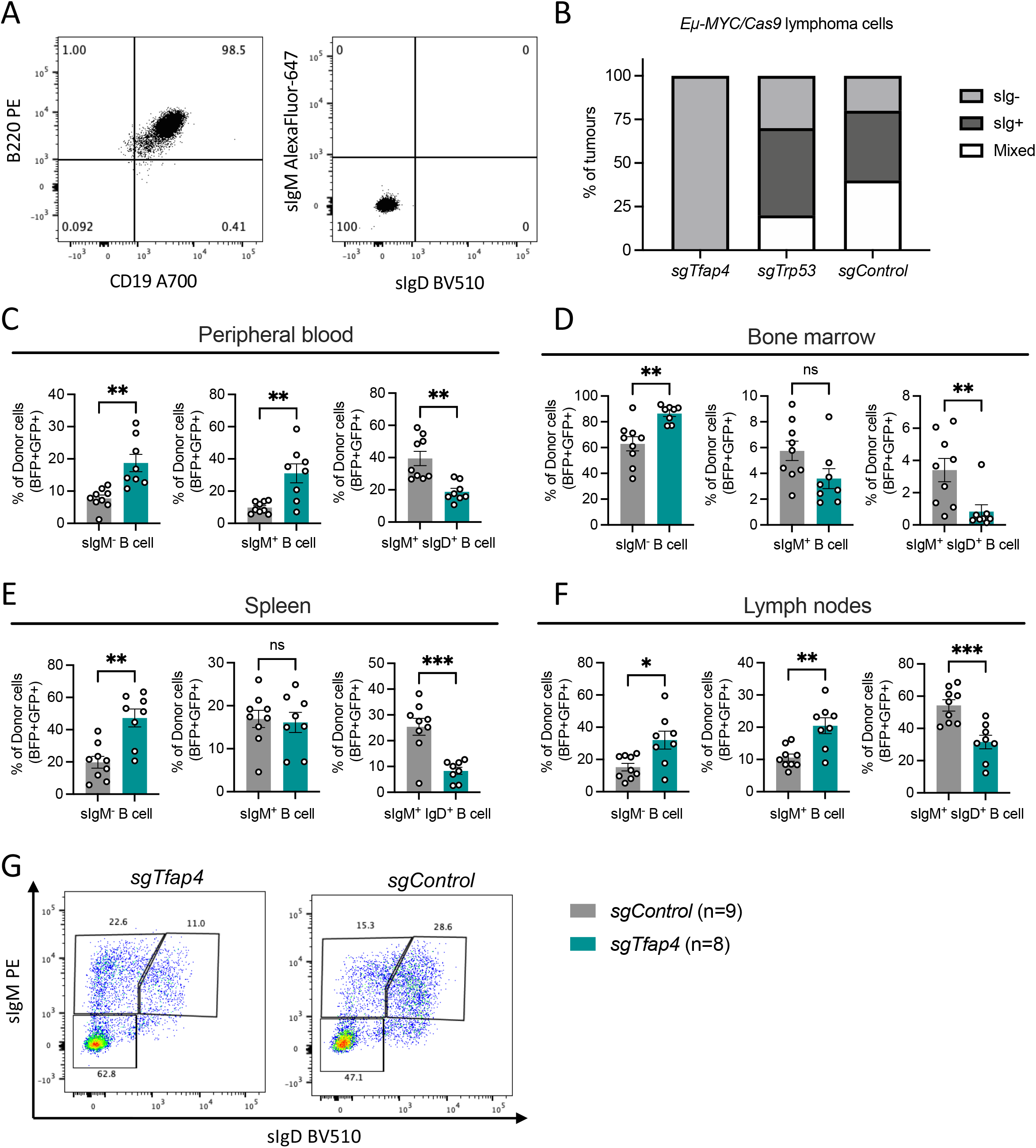
Deletion of TFAP4 in pre-leukemic *Eμ-MYC* pre-B cells impairs B cell differentiation. **A,** Representative flow cytometry plot of surface staining for B220, CD19, IgM and IgD on a *sgTfap4/Eμ-MYC/Cas9* lymphoma cell line. **B,** Summary graph of surface Ig staining by flow cytometry on cell lines derived from *Eμ-MYC/Cas9* primary lymphomas of each genotype; *sgTfap4*(n=10)*, sgTrp53* (n=20) and *sgControl* (n=5). sig^-^ represents B220^+^ slgM^-^ slgD^-^ lymphomas; slg^+^ represents B220^+^ sIgM^+^/sIgD^+^ lymphomas; and mixed indicates B220^+^ lymphoma cells with both sIgM^-^ and sIgM^+^ lymphoma cell populations. Lethally irradiated wild-type mice were reconstituted with *Eμ-MYC/Cas9* HSPCs transduced with either a *sgTfap4* or a *sgControl* vector. The hematopoietic cell subsets in these transplanted mice were analyzed at 3 weeks post-transplantation by flow cytometry. The percentages of donor derived (GFP^+^ BFP^+^) B cell subsets; sIgM^-^ (B220^+^ sIgM/sIgD^-^), immature sIgM^+^ B cells (B220^+^ sIgM^+^ sIgD^-^) and mature sIgM^+^/sIgD^+^ B cells (B220^+^ sIgM/sIgD^+^) in the peripheral blood (**C),** bone marrow **(D)**, spleen **(E)**, and lymph nodes **(F)** of recipient mice were determined. **G,** Representative flow cytometry dot plot to examine the different B cell subsets, gated on live donor derived lymphoid cells GFP^+^ BFP^+^ CD45.2^+^ B220^+^. Each dot represents an individual recipient mouse that had been transplanted with *sgTFAP4/Eμ-MYC/Cas9* (n = 8) or *sgControl/Eμ-MYC/Cas9* (n = 9) fetal liver cells, with error bars representing ±SEM. Unpaired two-tailed Student’s *t-*test with Welch’s correction, \**P*<0.05, **<0.01, ***<0.001.

Peripheral blood, bone marrow, spleen and lymph nodes showed significant differences in the B cell compartment between the *sgTfap4/Eμ-MYC/Cas9 vs* the *sgControl/Eμ-MYC/Cas9* HSPC transplanted cohorts. We detected higher proportions of surface Ig negative pro-B/pre-B cells (B220^+^ sIgM^-^ sIgD^-^) and a reduction in the proportions of mature B cells (B220^+^ sIg^+^) in the *sgTfap4/Eμ-MYC/Cas9* pre-leukemic recipient mice compared to their *sgControl/Eμ-MYC/Cas9* counterparts (Figure 3c-g). This accumulation of pro-B/pre-B cells in the *sgTfap4/Eμ-MYC/Cas9* pre-leukemic recipients is consistent with the immuno-phenotyping of the malignant lymphomas arising in these mice (Figure 3 a, b).

### TFAP4 deleted *Eμ-MYC* pre-B cells fail to upregulate transcriptional programs required for B cell differentiation

As TFAP4 is a transcriptional regulator, we next sought to understand how its loss might alter cell signaling pathways to accelerate c-MYC-driven lymphoma development. To this end we performed again a pre-leukemic analysis by transplanting *sgTfap4* or *sgControl* transduced *Eμ-MYC/Cas9* HSPCs into lethally irradiated wt mice. 21 days later we examined the B cell subsets in the bone marrow, the site of B cell development, by flow cytometry. We observed a significant reduction in the percentages of donor derived mature B cells (B220^+^sIgM^+^ sIgD^+^) and an increase in pro-B (B220^+^ sIgM^-^ cKIT^+^) and pre-B cells (B220^+^ sIgM^-^ cKIT^-^) in the *sgTfap4/Eμ-MYC/Cas9* recipient mice compared to the *sgControl/Eμ-MYC/Cas9* control mice (Figure 4a).

**Figure 4.**
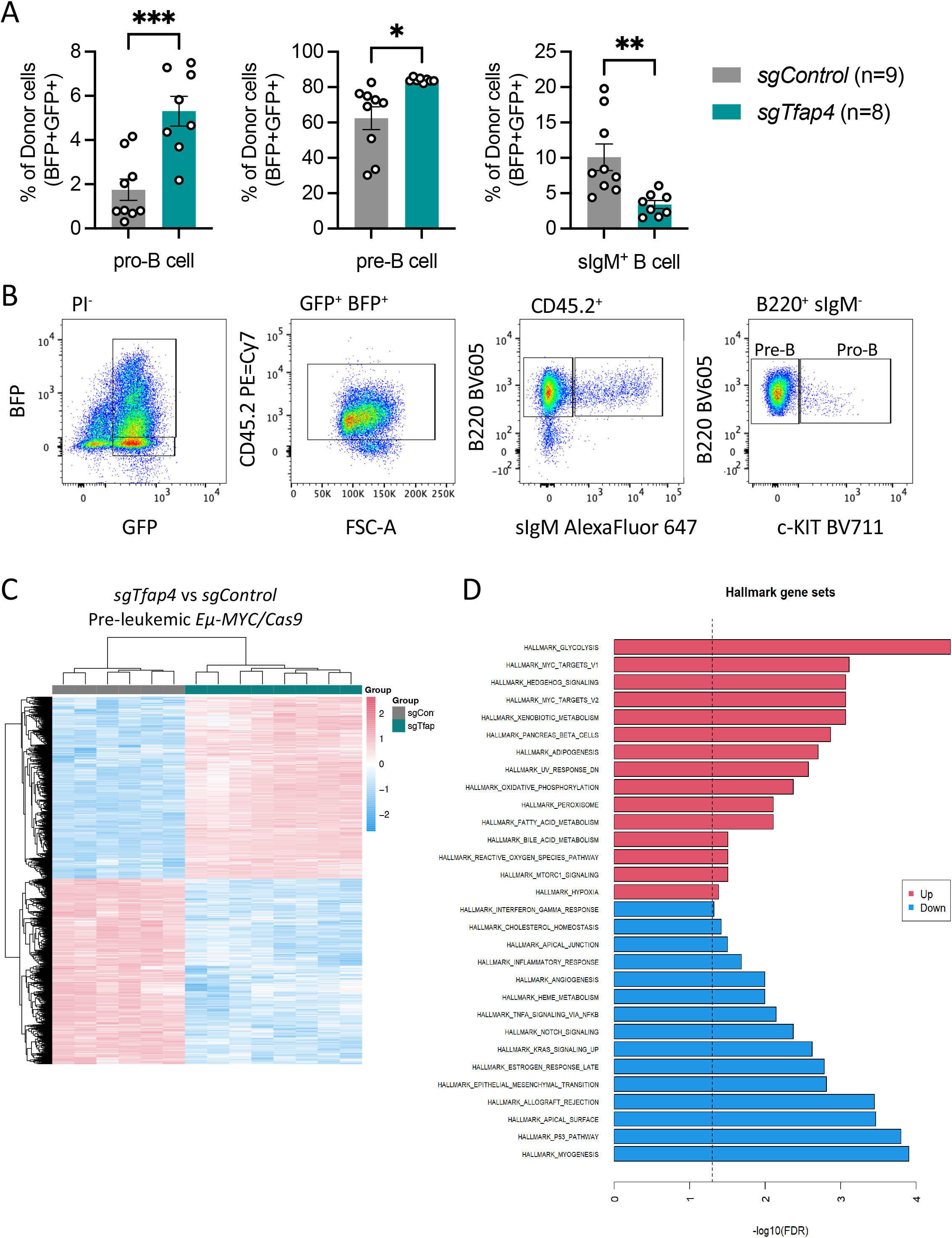
Pre-leukemic *Eμ-MYC/Cas9* pre-B cells lacking TFAP4 are transcriptionally distinct from control *Eμ-MYC/Cas9* pre-B cell. Lethally irradiated recipient mice were reconstituted with *Eμ-MYC/Cas9* fetal liver cells that had been transduced with either a vector encoding *sgTfap4* or a *sgControl*, and their pre-leukemic cells were analyzed at 3 weeks post-transplantation. **A,** In the bone marrow, the percentages of pro-B cells, pre-B cells and immature B cells derived from donor (GFP^+^ BFP^+^) HSPCs. Each dot represents an individual recipient mouse that had been transplanted with *sgTFAP4* (n = 8) or *sgControl* (n = 9) transduced *Eμ-MYC/Cas9* fetal liver cells, with error bars presenting mean ±SEM. Unpaired two-tailed Student’s *t-*test with Welch’s correction, \**P*<0.05, **<0.01, ***<0.001. **B,** Representative flow cytometry plots demonstrating gating strategy to identify the different donor derived (GFP^+^BFP^+^ CD45.2^+^) B cell subsets in the bone marrow of recipient mice: pro-B (B220^+^ sIgM^-^ c-KIT^+^), pre-B (B220^+^ sIgM^-^ c-KIT^-^), immature B (B220^+^ sIgM^+^) cells. **C-D,** RNA-seq analysis of donor derived pre-leukemic *Eμ-MYC/Cas9* pre-B cells (GFP^+^ BFP^+^ B220^+^ sIgM^-^ c-KIT^-^) from *sgTfap4/Eμ-MYC/Cas9* (n=8) or *sgControl/Eμ-MYC/Cas9* (n=9) cohorts isolated from the bone marrow of recipient mice. **C,** Heatmap showing differential gene expression between the pre-leukemic *sgTfap4l/Eμ-MYC/Cas9* pre-B cells and the *sgControl/Eμ-MYC/Cas9* control pre-leukemic pre-B cells. **D,** Hallmark gene set pathway analysis showing differentially expressed pathways between pre-leukemic *sgTfap4/Eμ-MYC/Cas9* pre-B cells *vs sgControl/Eμ-MYC/Cas9* pre-leukemic pre-B cells.

We isolated donor derived pre-leukemic pre-B cells (GFP^+^ BFP^+^ B220^+^ sIgM^-^ cKIT^-^) from both the *sgTfap4/Eμ-MYC/Cas9* recipient mice and the *sgControl/Eμ-MYC/Cas9* control counterparts (Figure 4b) and analyzed their transcriptional program by RNA-sequencing. Since fully transformed *Eμ-MYC* lymphomas are monoclonal they express only one Ig species. Our analysis revealed that the pre-leukemic *sgTfap4/Eμ-MYC/Cas9* and *sgControl/Eμ-MYC/Cas9* pre-B cells expressed a variety of Ig transcripts, suggesting that they were not fully transformed (Supplementary Figure 4a). The absence of TFAP4 markedly altered the transcriptional profile of the pre-leukemic *Eμ-MYC* pre-B cells (Figure 4c), with many hallmark gene set pathways differentially expressed in the pre-leukemic *sgTfap4/Eμ-MYC/* Cas9 pre-B cells compared to the *sgControl/Eμ-MYC/Cas9* control counterparts (Figure 4d).

3,218 genes were differentially expressed between the *sgTfap4/Eμ-MYC/Cas9 vs sgControl/Eμ-MYC/Cas9* pre-leukemic pre-B cells (Supplementary Figure 4b). To elucidate the contribution of loss of TFAP4 to c-MYC-driven lymphoma development we compared the expression of genes within certain hallmark cancer pathways. Consistent with the *in vitro* cell death assays using BH3 mimetic drugs targeting BCL-2 or MCL-1 in malignant *Eμ-MYC* lymphoma cells, we observed no differences in the expression of any of the genes for the pro-survival or pro-apoptotic BCL-2 family members (Figure 5a). Deregulated cell proliferation is also a hallmark of cancer (Hanahan, 2022) and mutations in TFAP4 have been implicated in abnormal cell proliferation in certain solid cancers (Wong *et al*., 2021). We observed no differences in the expression of *Mki67* (or Ki67), a marker of cell division, or cell cycle genes between the *sgTfap4/Eμ-MYC/Cas9 vs* the *sgControl/Eμ-MYC/Cas9* pre-leukemic pre-B cells (Figure 5b). This indicates that increased cell proliferation is not the mechanism by which loss of TFAP4 accelerates MYC-driven lymphoma development.

**Figure 5.**
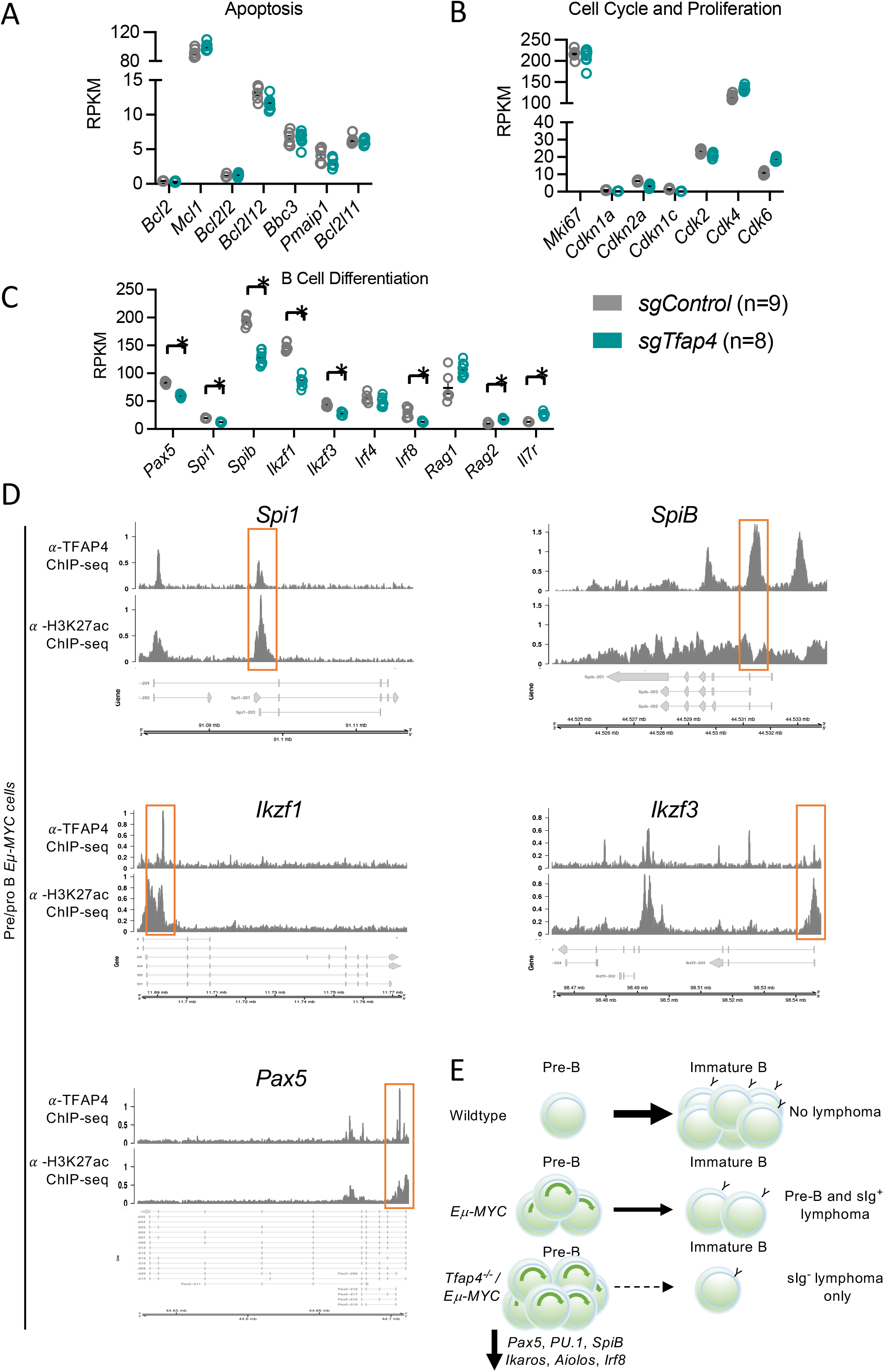
Genes regulating B cell differentiation are down-regulated in TFAP4 deleted pre-leukemic *Eμ-MYC* pre-B cells. Lethally irradiated recipient mice were transplanted with *Eμ-MYC/Cas9* HSPCs that had been transduced with a vector containing either a *sgTfap4* or a *sgControl*, and the pre-leukemic pre-B cells in these recipient mice were analyzed at 3 weeks post-transplantation by RNA-sequencing. **A-C,** Relative expression (RPKM – reads per kilobase million) of selected gene transcripts regulating apoptosis **(A)**, cell cycling and proliferation **(B)**, or master transcription factors and coordinators of B cell differentiation **(C).** Statistical significance *adj.*P* < 0.05, FDR < 0.05. **D,** *Spi1, SpiB, Ikzf1, Ikzf3* and *Pax5* genomic loci coverage plots of anti-TFAP4 and anti-H3K27ac (a marker of accessible chromatin) binding in wildtype pre-leukemic *Eμ-MYC* pro/pre-B cells. Y-axis represents counts per million (CPM), gene isoforms pictured, thick bar indicates exons, orange box highlights binding. **E,** Proposed mechanism: In wild-type mice normal B cell development occurs, whereby pre-B cells differentiate into immature B cells expressing surface IgM. In *Eμ-MYC* transgenic mice, c-MYC is abnormally over-expressed in the B cell lineage causing excess proliferation of pro-B/pre-B cells and a partial block in their differentiation, thereby producing an abnormally expanded pool of pre-leukemic pre-B cells and some sIg^+^ B cells. Some of these cells will acquire oncogenic mutations that can collaborate with c-MYC over-expression in neoplastic transformation and consequently give rise to pre-B or B cell lymphoma. The absence of TFAP4 in the *Eμ-MYC* setting results in an even larger pool of pre-leukemic pre-B cells arising due to the further restriction of differentiation by downregulation of transcription factors that are critical for B cell differentiation. This even larger pool of highly proliferative pre-leukemic pre-B cells is thus more likely to acquire additional mutations that drive transformation into malignant surface Ig^-^ pre-B cell lymphoma.

Our pre-leukemic analysis and the immuno-phenotyping of the malignant *Eμ-MYC* lymphoma cells revealed that the absence of TFAP4 results in the accumulation of pre-B cells, indicating a block in B cell development as a possible cause of accelerated c-MYC-driven lymphomagenesis. Therefore, we assessed the expression of genes encoding master transcriptional regulators of B cell differentiation. Interestingly, sg*Tfap4/Eμ-MYC/Cas9* pre-leukemic pre-B cells expressed lower levels of *Pax5*, the master regulator of B cell lineage specification (Figure 5c). Furthermore, we observed a reduction in *Spi1* (or PU.1), *SpiB, Irf8, Ikzf1* (Ikaros) and *Ikzf3* (Aiolos), genes that orchestrate differentiation of common lymphoid progenitors into pro-B, pre-B and finally immature B cells (Figure 5c) (Pang *et al*, 2014). Interestingly, TFAP4 ChIP-sequencing data in pre-leukemic *Eμ-MYC* pro-/pre-B cells demonstrates direct binding to *Ikzf1, Ikzf3, Spi1, SpiB* and *Pax5*(Figure 5d, Supplementary Figure 4c). These findings demonstrate that the absence of TFAP4 impairs the differentiation of highly proliferative *Eμ-MYC* pre-leukemic pre-B cells by reducing the levels of transcription factors that are critical for B cell differentiation (Figure 5e).

## Discussion

Here, we demonstrate that CRISPR/Cas9 mediated deletion of the transcription factor TFAP4 substantially accelerates c-MYC-driven pre-B lymphoma development. Flow cytometric analysis of fully transformed lymphoma cells and pre-leukemic *sgTfap4/Eμ-MYC/Cas9* B lymphoid cells revealed that the absence of TFAP4 imposes a block in B cell differentiation. No alterations in the TRP53 pathway or differences in expression of genes regulating apoptosis or cell cycling and proliferation were observed. However, aberrantly reduced expression of master transcriptional regulators of B cell differentiation, *Ikzf1, Ikzf3, Spi1, SpiB* and *Pax5*, direct target genes of TFAP4, was observed. This reveals that TFAP4 is required for the differentiation and maturation of *Eμ-MYC* pre-leukemic pre-B cells and thereby suppresses pre-B cell lymphoma development.

*Tfap4* is a direct c-MYC target gene, reported to contribute to certain c-MYC regulated cellular processes, including apoptosis, cell proliferation and cellular senescence (Jung & Hermeking, 2009). Induction of *Tfap4* by c-MYC to coordinate these processes has been implicated in the neoplastic transformation of several solid cancers (Wong *et al*., 2021). This contrasts our findings, where not over-expression but rather the absence of TFAP4 accelerated c-MYC-driven lymphomagenesis. In the context of c-MYC-driven lymphoma development, loss of TFAP4 does not alter apoptosis, cell cycling or cellular senescence as the genes regulating these processes were not differentially expressed in pre-leukemic pre-B cells lacking TFAP4. This indicates that the absence of TFAP4 promotes lymphoma development through an alternate mechanism, such as a block in B cell differentiation representing an emerging hallmark of cancer (Hanahan, 2022).

Recently, Tonc *et al.(Tonc et al., 2021*) showed that *Tfap4* is frequently mutated in human B cell malignancies. Additionally, they showed that loss of *Tfap4*, either a single allele or both alleles, accelerated c-MYC-driven lymphoma development in mice. This was done by employing both whole body *Tfap4* knockout and conditional *Tfap4* knockout mouse models crossed with *Eμ-MYC* transgenic mice as well as with other mouse models of c-MYC driven lymphoid malignancies. Concordant with our findings, all tumors Tonc *et al*., found in their *Tfap4^-/-^ Eμ-MYC* mice were pro-B/pre-B cell lymphomas. However, their pre-leukemic analysis using *Tfap4^+/-^ Eμ-MYC* mice showed no differences in the numbers of cells within the different B cell subsets (Tonc *et al*., 2021). The difference between their findings and our observations is likely due to the different model systems employed. In our case, the use of CRISPR/Cas9 results in complete deletion of *Tfap4*, evident by loss of TFAP4 protein (Figure 2a), whereas the pre-leukemic cells Tonc *et al*., examined still retained one allele of *Tfap4*, albeit some samples selected for the loss of the second allele, the latter highlighting that there is potent selection for loss of *Tfap4* in c-MYC-driven lymphoma development. Despite not observing differences in the proportions of B cell subsets in their pre-leukemic mice, Tonc *et al. (Tonc et al., 2021*) concluded that TFAP4 restricts c-MYC regulated B cell stemness because they found an upregulation of *Erg*, a direct c-MYC and TFAP4 target gene that regulates hematopoietic stemness. While we also observed increased levels of *Erg* in the *sgTfap4/Eμ-MYC/Cas9* pre-leukemic pre-B cells in our RNAseq analysis (data not shown), the major differences we found was a reduction in master regulators of B cell differentiation. We observed abnormally reduced expression of PU.1 (*Spi1), SpiB, Ikaros (Ikzf1*) and *Aiolos (Ikzf3*) in *sgTfap4/Eμ-MYC/Cas9* pre-leukemic pre-B cells; these are direct target genes of TFAP4, identified in pre-leukemic *Eμ-MYC* pre/pro-B cells. The contribution of these genes to normal B cell development and the impact of defects in their expression have been well characterized (reviewed by Pang *et al., (Pang et al., 2014)*). Interestingly, the deletion of PU.1 (Spi1) or SpiB impairs B cell differentiation, and mice lacking these transcriptional regulators develop B-ALL (Sokalski *et al*, 2011; Xu *et al*, 2012). Furthermore, loss of Ikaros (*Ikzf1*) or Aiolos (*Ikzf3*) is associated with B-ALL development in both mice and humans (Mullighan *et al*, 2007; Mullighan *et al*, 2009; Pang *et al*., 2014). *Pax5* deletion has been shown to prevent B cell differentiation beyond the pro-B cell stage (Pang *et al*., 2014) and its knockdown drives the development of B progenitor cell acute lymphoblastic leukemia (B-ALL) due to a differentiation block (Liu *et al*, 2014). We therefore propose that the absence of TFAP4 impairs the differentiation of c-MYC over-expressing pre-B cells through downregulation of PU.1, *Spi*B, *Ikaros, Aiolos* and *Pax5*. The reduced activation of these genes in the absence of TFAP4 prevents normal B cell differentiation causing an increase in the pool of highly proliferative pro-B/pre-B cells facilitating the acquisition of oncogenic lesions that cooperate with deregulated c-MYC expression in lymphomagenesis (Figure 5e). Thus, restoring TFAP4 expression might ameliorate disease progression, not by killing the tumour cells, but by inducing normal differentiation of the self-renewing pre-B cell lymphoma pool into less proliferative immature/mature B cells, albeit only if these cells are dependent on sustained absence of TFAP4 for lymphoma maintenance and not only for lymphoma development.

## Materials and methods

### sgRNA lentiviral vectors

Two independent constitutively expressed sgRNAs targeting *Tfap4* (sgRNA-1 5’-CGCATGCAGAGTATCAACGCGG; sgRNA-2 5’GATTGCCAACAGCAACGAGCGG) in a vector also containing a constitutively expressed BFP tag were obtained from the Merck CRISPR glycerol stock arrayed whole mouse genome sgRNA library, available at WEHI (www.sigmaaldrich.com MSANGERG).A positive control sgRNA targeting *Trp53* (5’-GGCAACTATGGCTTCCACCT); and a negative control sgRNA targeting human *BIM* (5’-GCCCAAGAGTTGCGGCGTAT), both in a vector also containing a BFP tag, were derived from the constitutive sgRNA FUGW expression vector previously described (Janic *et al*, 2018) and used for original hematopoietic reconstitution experiments. A vector constitutively expressing a negative control sgRNA targeting human *NLRC5* (sgRNA-1 5’-GCTGCAGAAGTGTCAGCTCCAGG) and the BFP tag was also obtained from the Merck CRISPR glycerol stock arrayed whole mouse genome sgRNA library, and this vector was used for the pre-leukemic analyzes.

### Cell culture and lentivirus production

HEK293T cells were maintained in DMEM (Gibco) medium containing 10% (v/v) heat inactivated fetal calf serum (HI-FCS; Gibco), 100 U/mL Penicillin (Sigma) and 100 μg/mL streptomycin (Sigma). *Eμ-MYC* lymphoma cell lines were derived from primary tumors and cultured in FMA medium - DMEM (Gibco) supplemented with 10% (v/v) HI-FCS (Gibco), 50 μM mercaptoethanol (Sigma-Alrich), 23.8 mM sodium bicarbonate (Merck), 1 mM HEPES (WEHI Media Kitchen), 13.5 μM folic acid (Sigma), 0.24 mM L-Asparagine monohydrate (Sigma), 0.55 mM L-Arginine monohydrochloride (Sigma), 22.2 mM D-glucose (Ajax), 100 U/mL Penicillin (Sigma) and 100 μg/mL streptomycin (Sigma) (as previously described; (Kelly *et al*., 2014)). Some *sgControl/Eμ-MYC/Cas9* lymphoma cell lines were previously used for experiments in (Mizutani *et al*., 2022). Lentiviruses were generated by transient transfection of HEK293T cells in 10 cm^2^ dishes using packaging the constructs pMDL (5 μg), RSV (2.5 μg) and pENV (5μg), and 10 μg of vector DNA by using standard calcium phosphate precipitation methods previously described (Herold *et al*, 2008). Supernatants containing lentivirus particles were collected 48-72 h after transfection and passed through a 0.45 μM filter prior to infection of cells.

### Experimental mice

The care and husbandry of experimental mice was conducted according to the guidelines outlined by The Walter and Eliza Hall Institute of Animal Ethics Committee. *Eμ-MYC* transgenic (Adams *et al*., 1985) and Cas9 transgenic mice (a gift from Prof K. Rajewsky, Max Delbrueck Centre, Berlin, Germany) were maintained on a C57BL/6-Ly5.2 background. C57BL/6-Ly5.1 and C57BL/6-Ly5.2 mice were obtained from the Walter and Eliza Hall Institute breeding facility (Kew, Victoria, Australia).

### Genotyping

Total DNA was extracted from ear-clips of adult mice or tails of day (E) 13.5 embryos using tail lysis buffer (Viagen Biotech) and proteinase K. DNA (1 μL) was PCR amplified in a mastermix of GoTaq Green (Promega) containing specific primers at a final concentration of 0.5 pmol/μL using the cycling conditions: 94°C for 4 min followed by 30 cycles of (94°C for 40 sec, 55°C for 30 sec, 72°C for 60 sec) and finally 72°C for 5 min. PCR products were size separated by gel electrophoresis on a gel composed of 2% DNA grade agarose (Bioline) in TAE buffer (40 mM Tris Acetate, 1 mM EDTA pH 8.0) containing ethidium bromide (0.2 μg/mL, Sigma) and imaged on GelDoc DOCTM XR+ Gel documentation system (Bio-Rad).

*Eμ-MYC:* ^~^900 bp
*MYC-1:* 5’ -CAGCTGGCGTAATAGCGAAGAG
*MYC-2:* 5’ -CTGTGACTGGTGAGTACTCAACC

### Hematopoietic reconstitution of lethally irradiated mice with fetal liver derived HSPCs

Fetal liver cells from *Eμ-MYC/Cas9* double transgenic day 13.5 embryos were harvested and frozen in 90% FCS and 10% (v/v) DMSO. Prior to transduction with sgRNA expression vectors, fetal liver cells were thawed and cultured for 48 h in alpha-minimum essential media (alpha-MEM) Glutamax (Gibco) containing 10% FCS, 1x Glutamax, 1 mM sodium pyruvate, 100 U/mL penicillin, 100 μg/mL streptomycin, 10 mM HEPES, 50 μM B-mercaptoethanol that was further supplemented with recombinant cytokines 10 ng/mL IL-6, 100 ng/mL mouse stem cell factor (SCF), 50 ng/mL thrombopoietin (TPO) and 10 ng/mL FLT-3 ligand kindly provided by Dr. Jian-Guo Zhang (WEHI). 12-well non-tissue culture treated plates were coated with 32 μg/mL retronectin (WEHI) in PBS solution overnight at 4°C then blocked with 2% bovine serum albumin solution (#A1595 Sigma-Aldrich) in PBS at 37°C for 30 min were prepared. Viral supernatant (produced as described above) supplemented with 8 μg/mL polybrene was centrifuged onto the retronectin coated plates at 3500 rpm for 2 h at 32°C. Supernatant was removed and replaced with fetal liver cells to be transduced for 24 h. These transduced cells were then collected and washed in PBS and intra-venously (i.v.) injected into lethally irradiated (2x 5.5 Gy, 4 h apart) C57BL/6-Ly5.1 mice. Transplanted mice were administered prophylactic neomycin dosed water and were carefully monitored for illness and euthanized by experienced animal technicians if they presented with weight loss, labored breathing, or palpable signs of lymphoma, including splenomegaly and lymphadenopathy. Tumour-free survival was defined as the time from HSPC transplantation until the animal developed lymphoma and had to be euthanized.

### Lymphoma analysis

Sick HSPC transplanted mice were euthanized and peripheral blood was collected and analyzed using an ADVIA hematology analyzer (Bayer). Hematopoietic tissues (spleen, thymus, lymph nodes – inguinal, axillary, brachial - and bone marrow from both femurs and tibiae) were harvested and mashed through 100 μM filters to generate single cell suspensions in PBS supplemented with 5% (v/v) FCS and 5 μM EDTA (pH 8.0) (hereafter called FACS buffer) and total cellularity was determined using an TC20 automated cell counter (Bio-Rad). Pellets of 2-5×10^6^ cells were collected for subsequent DNA and protein analysis. Lymphoma cell lines were derived by plating serial dilutions of tumour cells from enlarged hematopoietic tissues in FMA medium (described above). Lymphoma cell lines were immunophenotyped by staining with fluorochrome conjugated antibodies against B220 (RA3-6B2), IgM (5.1), IgD (11-26C), CD19 (ID3) and T cell antigen receptor beta (TCRβ) (H57-597) in FACS buffer supplemented with Fc block (2.4G2 supernatant, WEHI). Donor cells were identified as Ly5.2 positive, Cas9-eGFP and sgRNA CFP or BFP positive. Dead cells were excluded by gating on propidium iodide (PI) negative cells and analyzed in a Fortessax20 flow cytometer (Becton Dickinson). Data were analyzed with FlowJo™ analysis software.

### Western blotting

Total protein was isolated from cell pellets of lymphoma bearing tissues by lysis in RIPA buffer (50 mM Tris-HCl, 150 mM NaCl, 1% NP-40, 0.5% DOC, 0.1% SDS) supplemented with complete protease inhibitor (Roche). Protein concentration was determined using the Bradford Assay (Bio-Rad), 20 μg of protein was prepared in Laemmli buffer (0.25 M Tris.HCl pH 6.8, 40% glycerol, 0.8% SDS, 0.1% bromophenol blue, 10% mercaptoethanol), denatured at 100°C for 5 min, and size fractionated by gel electrophoresis on 4-12% NuPAGE™ Bis-Tris polyacrylamide gels (Thermo Fisher Scientific) in MES Running Buffer (50 mM MES - 4 morpholine ethane sulfonic acid, 50 mM Tris base, 1mM EDTA, 0.1% w/v SDS). Protein was then transferred onto nitrocellulose membranes (Life Technologies) using the iBlot membrane transfer system (Thermo Fisher Scientific) and blocked in PBS-T (PBS, 0.1% (v/v) Tween20) and 5% (w/v) skim milk. The following primary antibodies were diluted in PBS-T; mouse-anti-TRP53 (CM5, Novocastra™ Leica Biosystems); mouse-anti-p19/ARF (5.C3.1, Rockland); mouse-anti-AP4 (A-8, Santa Cruz Biotechnology, INC.); and mouse-anti-HSP70 (N-6, a gift from Dr. R. Anderson, Olivia Newton John Centre, Melbourne, Australia) served as a protein loading control. Secondary goat-anti-mouse/rat/rabbit IgG antibodies conjugated to HRP (Southern Biotech) were then applied. Luminata Forte Western horseradish peroxidase (HRP) substrate (Merck Millipore) was added prior to imaging using the ChemiDoc Imaging System with ImageLab software (Bio-Rad).

### Cell viability assays

5 x 10^4^ cells were plated in FMA medium (described above) in triplicate into 96-well flat-bottom plates and treated with serial dilutions of nutlin-3A (Cayman Chemical, #18585), etoposide (Ebewe Pharmaceuticals Ltd, A-4866), S63845 (Active Biochem, #6044) (Kotschy *et al*., 2016) or ABT-199/Venetoclax (Active Biochem, #A-1231) (Roberts *et al*., 2016). Cell viability was determined at 24 h by staining cells with 2 μg/mL PI (Sigma) and AnnexinV (AnnV) conjugated to Alexa Fluor 647 (WEHI) followed by analysis in a Fortessax20 flow cytometer (Becton Dickinson). Data were analyzed using FlowJo™ analysis software. Cell viability was normalized to DMSO vehicle control treated cells.

### Analysis of pre-leukemic pre-B cells

Reconstitution of lethally irradiated mice with fetal liver derived HSPCs was performed as described above, except fetal livers were not pooled prior to infection and instead one fetal liver was transduced and injected into two recipient mice, and recipient mice were harvested at three weeks (19 or 21 days) post-transplantation. From these mice, peripheral blood was collected by retro-orbital bleed into EDTA containing tubes and the spleen, thymus, lymph nodes (inguinal, brachial, axillary), and bone marrow (both femurs and tibiae) were harvested and processed into single cell suspensions and counted as described above. One to two million cells from each tissue and 10 μL of blood were collected. Red blood cells were removed from peripheral blood and spleen cell suspensions by incubation in red cell removal buffer (RCRB: 156 mM Ammonium Chloride, 11.9 mM Sodium Bicarbonate, 0.097mM EDTA) for 5 min on ice and then washed twice. Cell suspensions were incubated in combinations of fluorochrome-conjugated antibodies against the following surface markers: B220 (RA3-6B2), IgM (5.1), IgD (11-26c), CD4 (GK1.5), CD8 (53.6.7), MAC-1 (M1/70), GR-1 (RB6-8C5), TCRβ (H57-597) and CD45.2 (S450-15-2). Staining with PI was used for dead cell exclusion and cells were analyzed in a Fortessa1 flow cytometer (Becton Dickinson).

### Isolation of pre-leukemic pre-B cells for RNA-sequence analysis

Pre-B cells were isolated from the bone marrow of recipient mice three weeks post-transplantation with either *sgTfap4/Eμ-MYC/Cas9* or *sgControl/Eμ-MYC/Cas9* fetal liver cells. Bone marrow (both femurs and tibia) were flushed with FACS buffer to generate a single cell suspension. B cells were enriched by staining bone marrow cells in a 120 μL cocktail of biotinylated antibodies against TER119 (Ly76), MAC-1 (M1/70), GR-1 (RB6-8C5) for 20 min on ice, washed, then incubated with 20 μL MagniSort Streptavidin Negative Selection Beads (ThermoFisher Scientific) for 5 min at room temperature and placed on a magnet for 5 min to capture bead tagged cells. This was done to remove undesired erythroid and myeloid cells. The supernatant was transferred into a fresh tube, washed, and resuspended in 100 μL of a cocktail of fluorochrome conjugated antibodies against B220 (RA3-6B2), IgM (5.1), c-KIT (2B8) and 1×10^6^ live (PI negative) pre-B cells (B220^+^ sIgM^-^ c-KIT^-^) were FACS sorted, centrifuged, and resuspended in 700 μL QIAzol Lysis Reagent (Qiagen) and stored at −80°C. Total RNA was extracted using the QIAgen miRNeasy Mini Kit according to manufacturer’s instructions including optional on-column DNAase digestion. The mRNA libraries were prepared from an input of 100 ng of total RNA and indexed using the TruSeq RNA samples Prep Kit (illumina) according to manufacturer’s protocol. The samples were sequenced on an Illumina NextSeq using paired-end sequencing.

### RNA sequencing analysis

All samples were aligned to the mm10 build of the mouse genome using the Rsubread aligner (v 2.0.1) (Liao *et al*, 2019) with >98% of fragments (read pairs) were mapped to genome. Then all fragments overlapping mouse genes were summarized into counts using Rsubread’s featureCounts function. Genes were identified using Gencode annotation to the mm10 genome (v M25), >83% of mapped fragments were assigned to genes for all samples. Differential gene expression (DGE)analysis was then undertaken using the limma (Ritchie *et al*, 2015) (v 3.44.3) and edgeR (Robinson *et al*, 2009) (v 3.30.3) software packages.

Prior to analysis all gender specific genes – Xist and those unique to the Y-chromosome were removed to avoid gender biases and non-protein coding genes, immunoglobulin genes and those genes identified as ‘To be Experimentally Confirmed (TEC)’ were also removed. Expression bases filtering was then performed using edgeR’s filterByExpr function with default parameters. A total of 15,846 genes remained for downstream analysis. Compositional differences between the samples were then normalized using the trimmed mean of M-values (TMM) method (Robinson & Oshlack, 2010) then transformed to log_2_ counts per million (CPM). The correlation between samples transplanted with HSPCs from each fetal liver was calculated using limma’s duplicate correlation function (Smyth *et al*, 2005). Differential expression between the KO and control sample groups was then assessed using linear models which incorporated the aforementioned correlation, and robust empirical Bayes moderated t-statistics with a trended prior variance (robust limma-trend pipeline with duplicate correlation (Phipson *et al*, 2016). DGE was assessed relative to a fold change threshold of 1.2 using limma’s treat function (McCarthy & Smyth, 2009). The Benjamini and Hochberg method was used to control the false discovery rate (FDR) below 5%. A total of 3,218 genes were found to be significantly differentially expressed between the two samples.

Pathway analyzes of the Gene Ontology (GO) and Kyoto Encyclopedia of Genes and Genomes (KEGG) were conducted using limma’s goana and kegga functions, respectively. Analysis of the Molecular Signatures Database Hallmark gene sets was achieved using limma’s fry gene set test. The mean-difference plot was generated using limma’s plotMD function. All heatmaps were created using the pheatmap software package.

### ChIP sequencing analysis

The ChIP-seq data is available from Gene Expression Omnibus, accession number GSE133514 (Tonc *et al*., 2021). All FastQ files were downloaded and aligned to the mm10 build of the mouse genome using the Rsubread aligner (v 2.10.5) (Liao *et al*., 2019). At least 90% of reads were successfully mapped for all samples. Following alignment PCR duplicate reads were marked using Sambamba (v 0.6.6).

To generate the coverage plots, the reads aligning to each gene from both the ChIP-seq and RNA-seq were first read into R and the coverage calculated for each sample using the GenomicAlignments package (v 1.31.1). Duplicate reads were excluded for the ChIP-seq samples. The coverage for each sample was then divided by its respective library size multiplied by 1 million to generate counts per million (CPM). The mean was then taken of the replicate samples to give the average coverage for each experimental group. The coverage plots were then produced using the Gviz package (v 1.40.1).

### Statistical analysis

Statistical analysis was performed using GraphPad Prism software. Data is presented as mean ±standard error of the mean (SEM) unless otherwise stated. Mouse survival curves were analyzed using Mantel-Cox statistical test. Statistical significance of differences between two groups was determined using Student’s t-test, Welch’s correction was applied if data did not fit a normal model; ANOVA test was used for comparing multiple groups. *In vitro* cell death curves were all log transformed and fitted to a non-linear regression model and the IC50 derived using GraphPad Prism software.

## Acknowledgements

The authors thank the WEHI screening lab for providing arrayed glycerol stocks of sgRNAs; Dr S. Wilcox for running the NGS; Drs S. Nutt and A. Kallies for insightful discussions. We thank the WEHI Bioservices staff D. Fayle, G. Siciliano, C. Epifanio and C. Gatt for animal care and husbandry; T. Nikolaou for irradiation of mice and ADVIA services; the WEHI core facilities for help with Flow Cytometry and Bioinformatics support and our colleagues in the Herold, Strasser and Kelly laboratories. This work was supported by grants and fellowships from the Australian National Health and Medical Research Council (NHMRC) (Project Grants 1159658,1186575, 1143105 and 1145728 to MJH, to MJH and AS, Ideas Grants Program Grant 1113133 to AS Fellowship1020363 to AS, 1156095 to MJH), the Leukemia and Lymphoma Society of America (LLS SCOR 7015-18 to AS, MJH), the Cancer Council of Victoria (project grant 1147328 and 2021 Grant In Aid to MJH, 1052309 and Venture Grant to MJH and AS), Phenomics Australia (to AJK and MJH). MAP is supported by an Australian Government Research Training Program Scholarship provided by the Australian Commonwealth Government and the University of Melbourne. SM was supported by the Uehara Memorial Foundation and JSPS Grant-in-Aid for Research Activity Start-up (20K22854). This project was supported by operational infrastructure grants through the Australian Government Independent Research Institute Infrastructure Support Scheme (361646 and 9000220) and the Victorian State Government Operational Infrastructure Support Program.

## Contributions

AS, MJH conceived and designed the study. MAP performed and analyzed experiments, with contributions from SM, AJK, LT, MP. ALG and CLWS performed bioinformatic analysis of RNAseq data. MAP, AS, MJH wrote the manuscript. All authors edited and contributed to the manuscript.

## Statement of Data Availability

Data will be made available upon request.

## References (Harvard style)

Adams JM, Harris AW, Pinkert CA, Corcoran LM, Alexander WS, Cory S, Palmiter RD, Brinster RL (1985) The c-myc oncogene driven by immunoglobulin enhancers induces lymphoid malignancy in transgenic mice. Nature 318: 533–538

Aubrey BJ, Kelly GL, Kueh AJ, Brennan MS, O’Connor L, Milla L, Wilcox S, Tai L, Strasser A, Herold MJ (2015) An inducible lentiviral guide RNA platform enables the identification of tumor-essential genes and tumor-promoting mutations in vivo. Cell Rep 10: 1422–1432

Boboila S, Lopez G, Yu J, Banerjee D, Kadenhe-Chiweshe A, Connolly EP, Kandel JJ, Rajbhandari P, Silva JM, Califano A et al (2018) Transcription factor activating protein 4 is synthetically lethal and a master regulator of MYCN-amplified neuroblastoma. Oncogene 37: 5451–5465

Campbell KJ, Bath ML, Turner ML, Vandenberg CJ, Bouillet P, Metcalf D, Scott CL, Cory S (2010) Elevated Mcl-1 perturbs lymphopoiesis, promotes transformation of hematopoietic stem/progenitor cells, and enhances drug resistance. Blood 116: 3197–3207

Chen C, Cai Q, He W, Lam TB, Lin J, Zhao Y, Chen X, Gu P, Huang H, Xue M et al (2017) AP4 modulated by the PI3K/AKT pathway promotes prostate cancer proliferation and metastasis of prostate cancer via upregulating L-plastin. Cell Death & Disease 8: e3060–e3060

Chen S, Chiu SK (2015) AP4 activates cell migration and EMT mediated by p53 in MDA-MB-231 breast carcinoma cells. Mol Cell Biochem 407: 57–68

Chou C, Verbaro DJ, Tonc E, Holmgren M, Cella M, Colonna M, Bhattacharya D, Egawa T (2016) The Transcription Factor AP4 Mediates Resolution of Chronic Viral Infection through Amplification of Germinal Center B Cell Responses. Immunity 45: 570–582

Dang CV (2012) MYC on the path to cancer. Cell 149: 22–35

Eischen CM, Weber JD, Roussel MF, Sherr CJ, Cleveland JL (1999) Disruption of the ARF-Mdm2-p53 tumor suppressor pathway in Myc-induced lymphomagenesis. Genes Dev 13: 2658–2669

Grande BM, Gerhard DS, Jiang A, Griner NB, Abramson JS, Alexander TB, Allen H, Ayers LW, Bethony JM, Bhatia K et al (2019) Genome-wide discovery of somatic coding and noncoding mutations in pediatric endemic and sporadic Burkitt lymphoma. Blood 133: 1313–1324

Haluska FG, Finver S, Tsujimoto Y, Croce CM (1986) The t(8; 14) chromosomal translocation occurring in B-cell malignancies results from mistakes in V-D-J joining. Nature 324: 158–161

Hanahan D (2022) Hallmarks of Cancer: New Dimensions. Cancer Discovery 12: 31–46

Harris AW, Pinkert CA, Crawford M, Langdon WY, Brinster RL, Adams JM (1988) The E mu-myc transgenic mouse. A model for high-incidence spontaneous lymphoma and leukemia of early B cells. J Exp Med 167: 353–371

Herold MJ, van den Brandt J, Seibler J, Reichardt HM (2008) Inducible and reversible gene silencing by stable integration of an shRNA-encoding lentivirus in transgenic rats. Proc Natl Acad Sci U S A 105: 18507–18512

Hu BS, Zhao G, Yu HF, Chen K, Dong JH, Tan JW (2013) High expression of AP-4 predicts poor prognosis for hepatocellular carcinoma after curative hepatectomy. Tumour Biol 34: 271–276

Huang T, Chen QF, Chang BY, Shen LJ, Li W, Wu PH, Fan WJ (2019) TFAP4 Promotes Hepatocellular Carcinoma Invasion and Metastasis via Activating the PI3K/AKT Signaling Pathway. Dis Markers 2019: 7129214

Jaeckel S, Kaller M, Jackstadt R, Gotz U, Muller S, Boos S, Horst D, Jung P, Hermeking H (2018) Ap4 is rate limiting for intestinal tumor formation by controlling the homeostasis of intestinal stem cells. Nat Commun 9: 3573

Janic A, Valente LJ, Wakefield MJ, Di Stefano L, Milla L, Wilcox S, Yang H, Tai L, Vandenberg CJ, Kueh AJ et al (2018) DNA repair processes are critical mediators of p53-dependent tumor suppression. Nat Med 24: 947–953

Jung P, Hermeking H (2009) The c-MYC-AP4-p21 cascade. Cell Cycle 8: 982–989

Kaymaz Y, Oduor CI, Yu H, Otieno JA, Ong’echa JM, Moormann AM, Bailey JA (2017) Comprehensive Transcriptome and Mutational Profiling of Endemic Burkitt Lymphoma Reveals EBV Type-Specific Differences. Mol Cancer Res 15: 563–576

Kelly GL, Grabow S, Glaser SP, Fitzsimmons L, Aubrey BJ, Okamoto T, Valente LJ, Robati M, Tai L, Fairlie WD et al (2014) Targeting of MCL-1 kills MYC-driven mouse and human lymphomas even when they bear mutations in p53. Genes Dev 28: 58–70

Kotschy A, Szlavik Z, Murray J, Davidson J, Maragno AL, Le Toumelin-Braizat G, Chanrion M, Kelly GL, Gong JN, Moujalled DM et al (2016) The MCL1 inhibitor S63845 is tolerable and effective in diverse cancer models. Nature 538: 477–482

Langdon WY, Harris AW, Cory S, Adams JM (1986) The c-myc oncogene perturbs B lymphocyte development in E-mu-myc transgenic mice. Cell 47: 11–18

Liao Y, Smyth GK, Shi W (2019) The R package Rsubread is easier, faster, cheaper and better for alignment and quantification of RNA sequencing reads. Nucleic Acids Res 47: e47

Liu GJ, Cimmino L, Jude JG, Hu Y, Witkowski MT, McKenzie MD, Kartal-Kaess M, Best SA, Tuohey L, Liao Y et al (2014) Pax5 loss imposes a reversible differentiation block in B-progenitor acute lymphoblastic leukemia. Genes Dev 28: 1337–1350

Liu X, Zhang B, Guo Y, Liang Q, Wu C, Wu L, Tao K, Wang G, Chen J (2012) Down-regulation of AP-4 inhibits proliferation, induces cell cycle arrest and promotes apoptosis in human gastric cancer cells. PLoS One 7: e37096

López C, Kleinheinz K, Aukema SM, Rohde M, Bernhart SH, Hübschmann D, Wagener R, Toprak UH, Raimondi F, Kreuz M et al (2019) Genomic and transcriptomic changes complement each other in the pathogenesis of sporadic Burkitt lymphoma. Nature Communications 10: 1459

McCarthy DJ, Smyth GK (2009) Testing significance relative to a fold-change threshold is a TREAT. Bioinformatics 25: 765–771

Mermod N, Williams TJ, Tjian R (1988) Enhancer binding factors AP-4 and AP-1 act in concert to activate SV40 late transcription in vitro. Nature 332: 557–561

Mizutani S, Potts MA, Deng Y, Giner G, Diepstraten S, Kueh AJ, Pal M, Healey G, Tai L, Wilcox S et al (2022) Genome-wide <em>in vivo</em> CRISPR screens identify GATOR1 as a TP53 induced tumour suppressor. bioRxiv: 2022.2002.2016.480657

Mullighan CG, Goorha S, Radtke I, Miller CB, Coustan-Smith E, Dalton JD, Girtman K, Mathew S, Ma J, Pounds SB et al (2007) Genome-wide analysis of genetic alterations in acute lymphoblastic leukaemia. Nature 446: 758–764

Mullighan CG, Su X, Zhang J, Radtke I, Phillips LAA, Miller CB, Ma J, Liu W, Cheng C, Schulman BA et al (2009) Deletion of IKZF1 and Prognosis in Acute Lymphoblastic Leukemia. New England Journal of Medicine 360: 470–480

Pang SH, Carotta S, Nutt SL (2014) Transcriptional control of pre-B cell development and leukemia prevention. Curr Top Microbiol Immunol 381: 189–213

Phipson B, Lee S, Majewski IJ, Alexander WS, Smyth GK (2016) ROBUST HYPERPARAMETER ESTIMATION PROTECTS AGAINST HYPERVARIABLE GENES AND IMPROVES POWER TO DETECT DIFFERENTIAL EXPRESSION. Ann Appl Stat 10: 946–963

Ritchie ME, Phipson B, Wu D, Hu Y, Law CW, Shi W, Smyth GK (2015) limma powers differential expression analyses for RNA-sequencing and microarray studies. Nucleic Acids Res 43: e47

Roberts AW, Davids MS, Pagel JM, Kahl BS, Puvvada SD, Gerecitano JF, Kipps TJ, Anderson MA, Brown JR, Gressick L et al (2016) Targeting BCL2 with Venetoclax in Relapsed Chronic Lymphocytic Leukemia. N Engl J Med 374: 311–322

Robinson MD, McCarthy DJ, Smyth GK (2009) edgeR: a Bioconductor package for differential expression analysis of digital gene expression data. Bioinformatics 26: 139–140

Robinson MD, Oshlack A (2010) A scaling normalization method for differential expression analysis of RNA-seq data. Genome Biology 11: R25

Shi L, Jackstadt R, Siemens H, Li H, Kirchner T, Hermeking H (2014) p53-induced miR-15a/16-1 and AP4 form a double-negative feedback loop to regulate epithelial-mesenchymal transition and metastasis in colorectal cancer. Cancer Res 74: 532–542

Smyth GK, Michaud J, Scott HS (2005) Use of within-array replicate spots for assessing differential expression in microarray experiments. Bioinformatics 21: 2067–2075

Sokalski KM, Li SKH, Welch I, Cadieux-Pitre H-AT, Gruca MR, DeKoter RP (2011) Deletion of genes encoding PU.1 and Spi-B in B cells impairs differentiation and induces pre-B cell acute lymphoblastic leukemia. Blood 118: 2801–2808

Strasser A, Harris AW, Bath ML, Cory S (1990) Novel primitive lymphoid tumours induced in transgenic mice by cooperation between myc and bcl-2. Nature 348: 331–333

Strasser A, Harris AW, Jacks T, Cory S (1994) DNA damage can induce apoptosis in proliferating lymphoid cells via p53-independent mechanisms inhibitable by Bcl-2. Cell 79: 329–339

Tonc E, Takeuchi Y, Chou C, Xia Y, Holmgren M, Fujii C, Raju S, Chang GS, Iwamoto M, Egawa T (2021) Unexpected suppression of tumorigenesis by c-MYC via TFAP4-dependent restriction of stemness in B lymphocytes. Blood 138: 2526–2538

Vassilev LT, Vu BT, Graves B, Carvajal D, Podlaski F, Filipovic Z, Kong N, Kammlott U, Lukacs C, Klein C et al (2004) In vivo activation of the p53 pathway by small-molecule antagonists of MDM2. Science 303: 844–848

Wei J, Yang P, Zhang T, Chen Z, Chen W, Wanglin L, He F, Wei F, Huang D, Zhong J et al (2017) Overexpression of transcription factor activating enhancer binding protein 4 (TFAP4) predicts poor prognosis for colorectal cancer patients. Exp Ther Med 14: 3057–3061

Wong MM, Joyson SM, Hermeking H, Chiu SK (2021) Transcription Factor AP4 Mediates Cell Fate Decisions: To Divide, Age, or Die. Cancers (Basel) 13

Xu LS, Sokalski KM, Hotke K, Christie DA, Zarnett O, Piskorz J, Thillainadesan G, Torchia J, DeKoter RP (2012) Regulation of B Cell Linker Protein Transcription by PU.1 and Spi-B in Murine B Cell Acute Lymphoblastic Leukemia. The Journal of Immunology 189: 3347–3354

Xue C, Yu DM, Gherardi S, Koach J, Milazzo G, Gamble L, Liu B, Valli E, Russell AJ, London WB et al (2016) MYCN promotes neuroblastoma malignancy by establishing a regulatory circuit with transcription factor AP4. Oncotarget 7: 54937–54951

